# A modified two-color flow cytometry assay to quantify in-vitro reinvasion and determine invasion phenotypes at low *Plasmodium falciparum* parasitemia

**DOI:** 10.1101/2019.12.20.885301

**Authors:** Ngoh Ines Atuh, Anong Damian Nota, Fru Jerome Cho, Fatoumata Bojang, Haddijatou Mbye, Alfred Amambua-Ngwa

## Abstract

Two-color flow cytometry(2cFCM) is the most accessible method for phenotyping parasite invasion. However, current protocols require samples of field isolates at ∼1% parasitemia for assay set-up, which are becoming more uncommon in low transmission settings. Current protocols, therefore, have to be adapted for low parasitemia if the method must have continued applicability in this era of elimination. Optimizing the protocol requires addressing; interference from young uninfected RBCs background fluorescence and biased phenotypes due to limited labeled RBCs availability and/or parasite density per assay. Here, we used SYBR Green I and CTFR Proliferation fluorescent dyes to set-up invasion assays with *Plasmodium falciparum* 3D7, Dd2 and field isolates cultures (diluted at 0.05% to 2.0% parasitemia) against varying unlabeled to labeled RBC ratios (1:1 to 1:4). We showed that a shorter SYBR Green I staining time of 20 minutes, down from 1hour, minimized background fluorescence from uninfected RBCs (mean= 0.03% events) and allowed 2cFCM to accurately quantify reinvasion for an assay at 0.02% parasitemia. An increase in labeled target RBCs to 1:3 per assays significantly increased heterologous reinvasion (p<0.001). This resulted in a 10% greater invasion inhibition by enzyme treatments (p<0.05). Strain-specific invasion phenotype could be accurately determined for samples with as low as 0.3% parasitemia. Samples above 0.8% parasitemia were less accurate. These findings show that invasion pathway phenotypes can be obtained for field samples with low parasitemia at greater sensitivity and reproducibility by increasing the proportion of labeled RBCs per assay by at least 2-fold what is in current methods.

## BACKGROUND

Invasion of human red blood cells (RBCs; Erythrocytes) by *Plasmodium falciparum* merozoite is vital for malaria parasite multiplication and transmission between hosts and it is considered a key target for the development of malaria interventions (Burns et al. 2019). Invasion mechanism makes use of multiple and redundant parasite ligand and erythrocyte receptor combinations (Perkins and Holt 1988; Binks and Conway 1999), whose dynamics are affected by host, transmission rates and environmental factors such as interventions. Studies on parasite invasion mechanisms and invasion phenotypes could, therefore, help in monitoring the effects of interventions and in designing alternative intervention strategies against natural parasite populations (Stubbs et al. 2005; Beeson et al. 2019). Such studies have recently been scaled up by the application of sensitive high throughput two-color Flow Cytometry (2cFCM)-based methods (Theron et al. 2010; Bei et al. 2010; Bei and Duraisingh 2015). These methods quantify the total number of successfully invaded merozoites (reinvasion), rather than just the total number of parasitized erythrocytes (growth). As such, quantifying reinvasion has wide applications in phenotyping parasites where it is used as a surrogate indicator of parasite viability or parasite multiplication in the presence of various stressors (Witkowski et al. 2013; Amaratunga et al. 2014; Douglas et al. 2011; Gomez-Escobar et al. 2010; Bowyer et al. 2015).

2cFCM is the preferred approach for phenotyping invasion pathways as it maintains RBC integrity and parasite viability. Unlike previous methods (Reed et al. 2000; Duraisingh et al. 2003; Jennings et al. 2007; Lantos et al. 2009), it does not require isolation and pre-enzyme treatment of infected donor RBCs so as to limit invasion of uninfected donor RBCs (autologous reinvasion). Conventional 2cFCM combines equal (1:1) proportions of infected unlabeled donor RBCs with uninfected fluorescently labeled target RBCs (pre-treated with neuraminidase, trypsin and chymotrypsin to remove various surface receptors) to specifically quantify parasite reinvasion into target RBCs (heterologous reinvasion) over a single erythrocytic cycle. Parasitized target RBCs are then detected and counted by flow cytometry following parasite nucleic acid staining with dyes such as SYBR Green I (Theron et al. 2010; Bei and Duraisingh 2015). However, current protocols are optimized for testing malaria samples with ∼1% parasitemia (pct), a level becoming less common in low transmission settings following massive interventions for malaria elimination.

For wider and continuous applicability of 2cFCM in this era of elimination, the current protocol needs to be adapted for increased sensitivity at low parasitemia. This will require; (a) reducing the high background fluorescence from nucleic acid-containing uninfected (unRBCs) by nucleic acid staining dyes like SYBR Green I (Theron et al. 2010), (b) minimizing autologous reinvasion, which can potentially create a bias in the measured phenotype. Autologous reinvasion is high with the high proportion of unlabeled donor unRBCs present in 1:1 ratio protocols (Bei and Duraisingh 2015). (c) determining the lower threshold of parasitemia for invasion phenotyping given that parasite invasion is highly strain-specific and parasite density-dependent (Rovira-Graells et al. 2016). Assays with too high or too low initial sample parasitemia could cause phenotype masking. This study, therefore, optimized the current 2cFCM protocol, adapting it to enable its application at low sample parasitemia in settings with declining malaria. This will allow for continuous monitoring of invasion phenotypes with the drive to elimination and evaluation of invasion-targeting candidate tools in low transmission settings.

## METHODS

### Parasite Strains and Cultures

*Plasmodium falciparum* 3D7 and Dd2 laboratory-adapted strains and three field isolates CMN45, CMN23 and CMN40 collected from the North-West region of Cameroon, were grown in O+ erythrocytes from a malaria-free adult donor at 2% hematocrit (HCT) in incomplete RPMI-1640 medium (iRPMI) supplemented with 0.5% albumax. Culture flasks were placed in an enclosed culture chamber and gassed with 1% O_2_, 5% CO_2_, and 94% N_2_. The chamber was then placed in a modular incubator unit (Billups-Rothenberg) at 37°C.

### Enzymatic Treatment of Target RBCs

Freshly washed O^+^ erythrocytes were treated with either 13μl of 33.35mU/ml Neuraminidase (Nm) (2UN/ml Sigma, Cat N6514), 60μl of 1.0mg/ml of Trypsin (high Trp) (sigma, Cat T9935) and 60μl of 1.0mg/ml of Chymotrypsin (high Chy) (sigma, Cat C4129) for 1 hour in separate tubes (with corresponding enzyme names; Nm, high Trp and high Chy) and incubated in a rotating chamber at 37°C. A positive control tube with all enzymes combined (Nm + Chy + Trp) and negative control with no enzyme (RPMI) were included. After incubation, the enzyme-treated RBCs were washed three times with 1X PBS and resuspended to 50% HCT in iRPMI and stored at 4°C for up to 24 hours before labelling. Efficacy of enzymatic treatment was confirmed by inhibition of invasion efficiency into Nm treated RBCs of <35% and >70% for 3D7 and Dd2 strains at 0.5% pct respectively as reported elsewhere (Bowyer et al. 2015).

### Labeling of Target RBCs

Target RBCs were fluorescently labeled with Cell Trace^(R)^ Far Red (CTFR) Proliferation Kit (Invitrogen C34564) at 2% HCT. The dye belongs to the same CTFR family of fluorophores as DDAO-SE (7-hydroxy-9H-(1, 3-dichloro-9, 9-dimethylacridin-2-one) succinimidyl ester. Due to its first-time use, optimization experiments for dye concentration (5 µM, 7.5 µM and 10 µM) and staining duration (30minutes, 1hr and 2hrs) were performed. RBCs were incubated at 37°C on a rotating wheel and washed three times after incubation. Stained RBCs were suspended at 2% HCT in cRPMI and stored for up to one week at 4°C for use in invasion assays.

### *In-vitro* invasion assay to determining the effect of parasite density and increasing labeled RBC proportion on 2cFCM sensitivity to quantify reinvasion at low parasitemia

The ratio of unlabeled donor to labeled target RBCs in assays was varied from 1:1 to 1:4 across four wells per ratio for each level of initial sample parasitemia. Briefly, stock infected donor cultures were synchronized to ∼90% ring stages with 5% w/v D-Sorbitol and diluted to a range of initial sample parasitemia of 0.05% to 2.0% by addition of O+ unRBCs. For each invasion assay, 20μl of infected culture (for each test parasitemia) was added to 50μl, 67μl, 75μl and 80μl of labeled target RBCs, corresponding to assay ratios of 1:1, 1:2, 1:3 and 1:4 respectively. The final reaction volume of 100μl at 2%HCT was constituted by adding 30μl, 13μl, 5μl and 0μl of unlabeled uninfected O^+^ RBCs respectively (figure 1). The assay parasitemia ranged from 0.02% to 0.2% in the wells. Parasitemia and developmental stage for each assay was determined by light microscopy of 10% Giemsa stained thin films. The average of two readers was retained. For invasion-inhibition assays with enzyme-treated RBCs, 20μl unlabeled iRBCs were added to 50μl and 75μl labeled enzyme-treated RBCs corresponding to assay ratios of 1:1 and 1:3 respectively. The final reaction volume of 100μl at 2%HCT was constituted by adding 30μl, and 5μl volumes of unlabeled Nm+Try+Chy treated RBCs. Triplicate well assays per sample were run and plates were incubated over a single erythrocytic cycle in flat-bottom 96 well plates (FALCON, USA). Isolates were cultured for ∼58 hours until reinvasion with the reemergence of post-8 hours young rings (figure 1).

**Figure 1.**
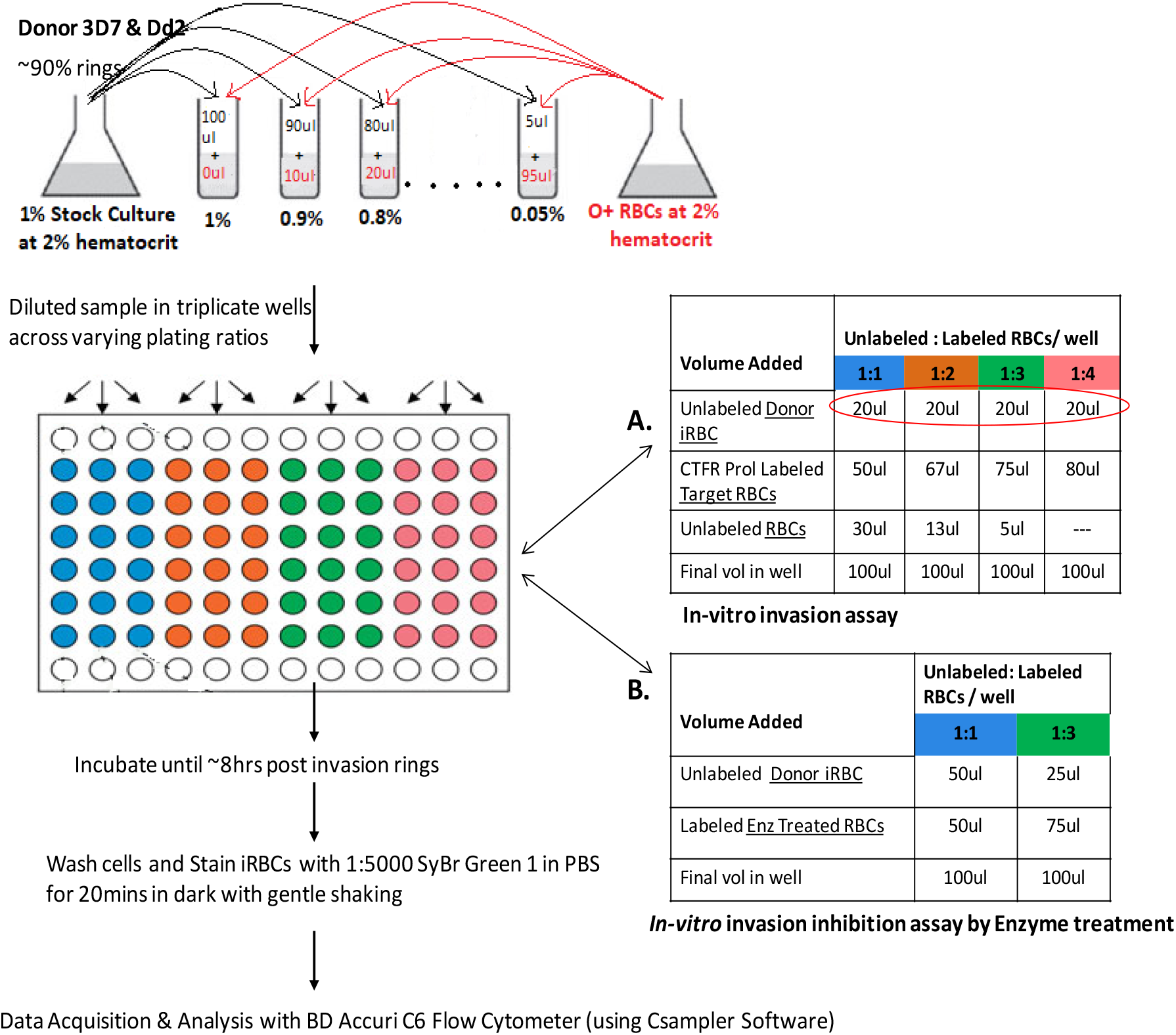
Experimental Protocol. Assessing the effect of parasite density and increased labeled RBC proportions per assay on the sensitivity of two-color FCM to quantify reinvasion and determine invasion pathways at low parasitemia.

The limit of quantification was determined as the minimum reinvasion parasitemia that could be detected and quantified for a corresponding starting parasitemia and cell ratio, after deducting the average background fluorescence for unRBCs.

### Flow cytometry data acquisition and analysis

Cell counting by flow cytometry was done with a BD Accuri™ C6 flow cytometer (BD Biosciences, Oxford, UK) equipped with a 488 nm 20 mW blue laser and a 633 nm 17 mW red laser for exciting SYBR Green1 (Invitrogen S7563) and CTFR Proliferation (Invitrogen C34564) dyes respectively. Parasite DNA in erythrocytes were detected following staining with SYBR Green I diluted in PBS to 1:5000 and incubated for 20minutes with gentle shaking in the dark. Detectors of forward scatter-FSC, side scatter-SSC, FL1 (SYBR Green I) and FL4 (CTFR Prol) were set in logarithmic scale and 100,000 events were collected for each assay (microplate well). Intact RBCs were selected with a polygonal gate on an FSC/SSC plot. Labelled target RBCs were gated on dot plots of SSC-H versus CTFR intensity (FL4-H) while iRBC were gated on dot plots of SSC-H versus SYBR Green I intensity (FL1-H). The cluster (gates) of iRBCs population was set at 10 fluorescent units away from unRBCs population to exclude weak fluorescence of RNA-containing uninfected young RBCs (figure 3A 1), previously localized with pico-green (Jun et al. 2012). Initial data visualization and analysis were performed using BD6 Csampler software.

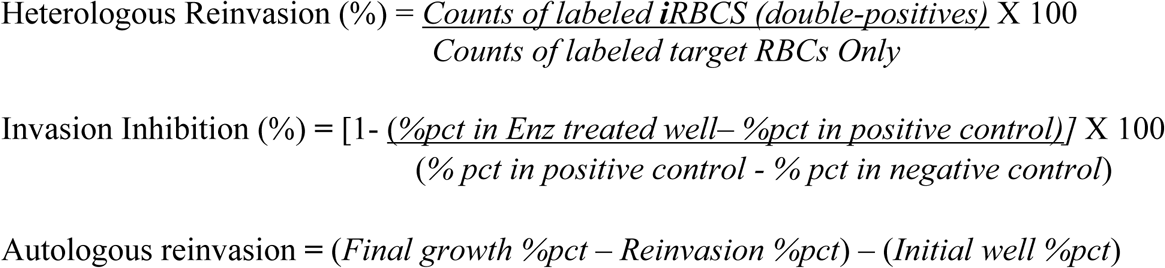

**Figure 2.**
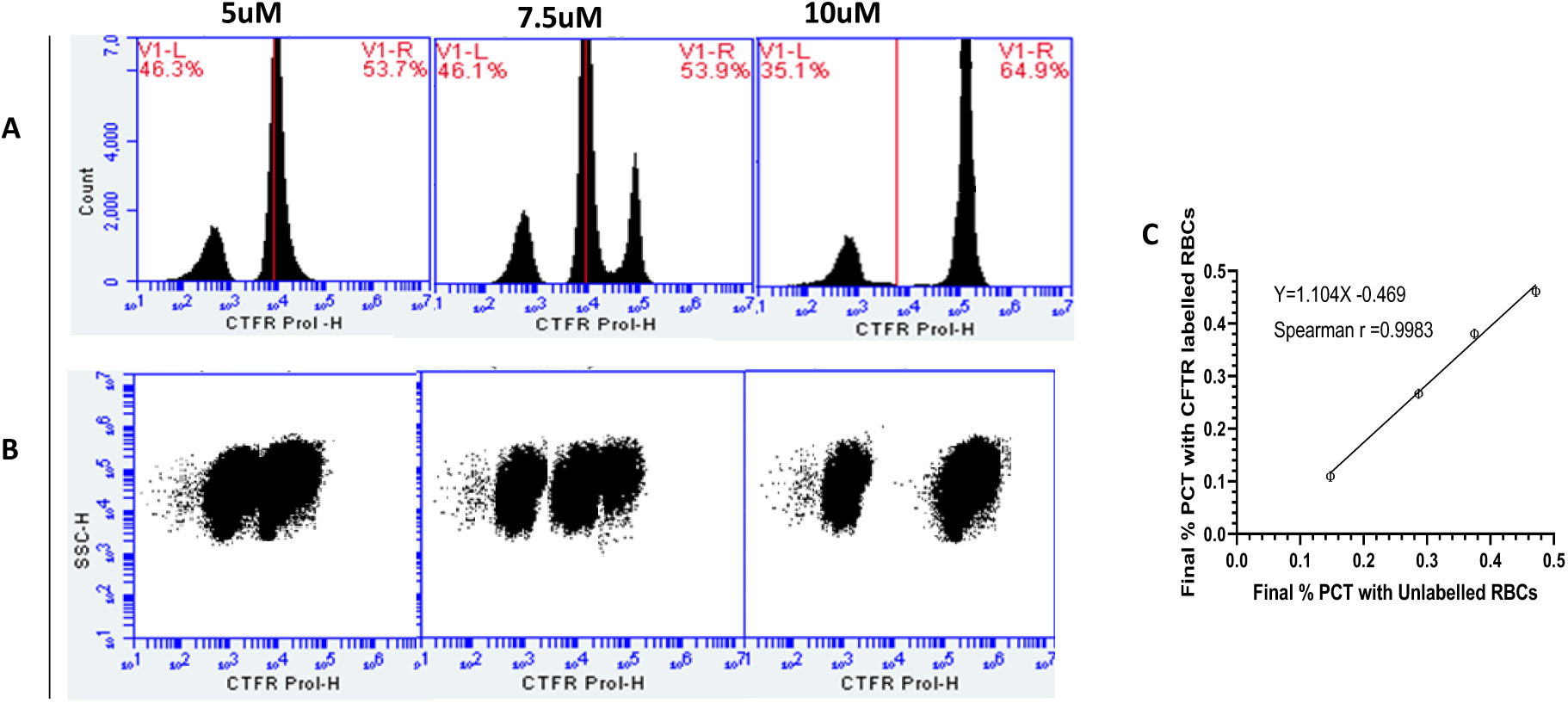
Labeling target RBCs with CTFR Proliferation Kit Dye for two-color FCM-based Invasion Assays. Target RBCs population labeled with Cell Trace Far Red (CTFR) Proliferation at 5uM, 7.5uM and 10uM concentrations for 1 hour. Labelled RBCs were co-incubated with donor unlabeled RBCs over a single erythrocytic cycle. (A) FCM discrimination of both RBC populations in histogram plots and (B) in dot plots. The dye also had no significant impact on parasite invasion efficiency. Final parasitemia of donor cultures (at 0.05%, 0.1%, 0.15% and 0.2% initial parasitemia) co-incubated with equal volumes of unlabeled RBCs and target labeled RBCs were highly correlated, Spearman *r =* 0.9963 (C).

**Figure 3.**
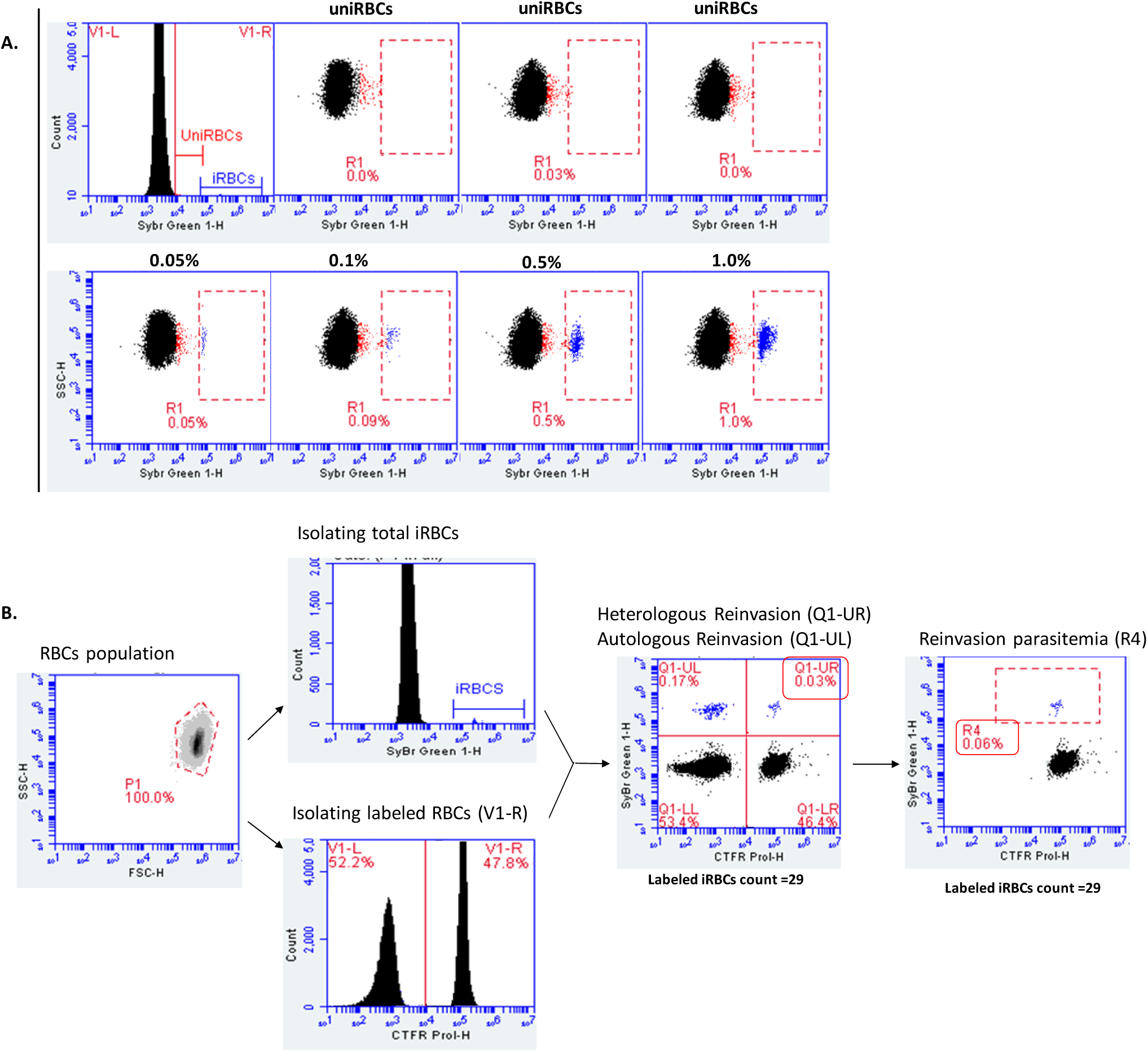
Accuracy of FCM enumeration of low parasitemia with SYBY Green 1 staining. (A) FCM determined parasitemia for triplicate uninfected RBC (uniRBCs) cultures in upper panel and 3D7 cultures at estimated parasitemia of 0.05%, 0.1%, 0.5% and 1.0% in the lower panel. Infected RBCs were stained with SYBR Green 1 diluted in PBS to 1:5000 for 20mins without RNase treatment. Estimated sample parasitemia was obtained from the microscopy determined value and the dilution factor. Gates for infected RBCs fluorescence was set at 10 units away from uninfected RBCs population to exclude the weak fluorescence from uninfected young RNA-containing RBCs (previously so localized with pico-green [21]). (B). Gating strategy for double positives (SYBR Green 1 + CTFR Prol. positive RBCs) for improved sensitivity.

“Double-positives” are the counts of RBCs positive for SYBR Green I and CTFR; positive control is Nm+Try+Chy treated and negative control is non enzyme-treated; final %pct is %pct of labelled target + unlabelled donor iRBCs, reinvasion %pct is %pct in labelled target RBCs and initial well %pct is diluted %pct in the well at assay set-up.

### Statistical Analysis

Plots and statistical analysis were made with Graph Pad prism vs 8.01. All data were normally distributed (passed Kolmogorov Smirnoff normality test at p=0.05) and were analyzed using parametric tests. A multiple t-test comparison for unpaired data was used for overall comparisons of % reinvasion and % invasion inhibition between plating ratios while a two-way Analysis of variance (ANOVA) with Bonferroni correction post hoc was used for subsequent pairwise comparisons of percent reinvasion and invasion inhibition for individual starting parasitemia within and between plating ratios.

## RESULTS

### Optimal concentration of CTFR Proliferation for Labeling Target RBCs

On histogram plots of cell counts, peaks of CTFR-labeled RBCs populations were readily detected after 1hr of incubation with the 5µM, 7.5µM and 10µM dye concentrations tested, with fluorescence intensity increasing with dye concentration (figure 2A). When these labeled RBCs were co-incubated with unlabeled infected RBCs over a single erythrocytic cycle, 10µM gave the highest resolution of the two RBC populations allowing for the best separation of donor from target infected RBCs in scatter plots (figure 2B). Hence 10µM of CTFR was chosen for all subsequent experiments. The dye also had no significant effect (*p=0*.*083*) on invasion efficiency as donor cultures at 0.05%, 0.1%, 0.15% and 0.2% parasitemia, co-incubated with equal volumes of target unlabeled and target labeled RBCs in separate assays gave highly correlated final parasitemia, spearman (r) = 0.9963 (figure 2C).

### Accuracy of FCM enumeration of low parasitemia with SYBR Green 1 staining

Parasite DNA was detected by staining iRBCs with SYBR Green 1 diluted in PBS to 1:5000 for 20minutes. SYBR Green 1 background fluorescence from unRBCs was very minimal with clusters (gates) of iRBCs population set at 10 fluorescent units away from unRBCs population to exclude weak fluorescence of RNA-containing uninfected young RBCs (previously so localized with pico-green (Jun et al. 2012)) (figure 3A). Background fluorescence varied from 0.0% to 0.03% of events for each triplicate assays (figure 3A upper panel). With this gating FCM counts of infected cells in *P. falciparum* 3D7 cultures diluted at 0.05%, 0.1%, 0.5% and 1.0% parasitemia was highly reproducible and determined at 0.05%, 0.09%, 0.5% and 1.0% respectively (figure 3A lower panel). From an assay with 0.1% starting parasitemia, reinvasion parasitemia increased from 0.03% (when double-positives were counted and reported as a percentage of the total RBCs population) to 0.06% (when double positives were counted and reported as a percentage of the target labeled RBC population only) (figure 3B).

### The effect of higher unlabeled to labeled RBCs ratios on the sensitivity of 2cFCM to quantify reinvasion at low parasitemia

FCM was accurate in quantifying reinvasion into labeled RBCs (heterologous reinvasion) down to 0.02% pct per assay (microplate well). At this parasitemia, reinvasion rate increased from 0.05% to 0.07%, 0.11% and 0.12% for 1:1, 1:2, 1:3 and 1:4 unlabeled to labeled RBCs per assay respectively. The increase in reinvasion rate with increasing labeled RBCs proportion was seen across the entire parasitemia range of 0.02% to 0.2% pct. When compared to 1:1 assays, the difference in the mean reinvasion parasitemia vary between *p*>*0*.*052* for 1:2 and *p*<*0*.*001* for 1:3 and 1:4 assays (figure 4A). On the other hand, reinvasion into unlabeled RBCs (autologous reinvasion) was significantly reduced with higher ratios (*p*>*0*.*05* for 1:2 and *p*<*0*.*001* for 1:3 and 1:4 assays). The magnitude of the reduction was at least five times in the 1:4 ratio. While autologous reinvasion values showed a steady increase with increasing parasite density in the 1:1 and 1:2 assays, values were very similar in 1:3 and 1:4 assays across the parasitemia range tested (figure 4B). Thus, reinvasion parasitemia (percentage of parasitized labeled RBCs) approximated the final growth parasitemia (percentage of parasitized labeled + unlabeled RBCs) when the labeled RBCs proportion in the well was two times higher as can be seen in supplementary figure.

**Figure 4.**
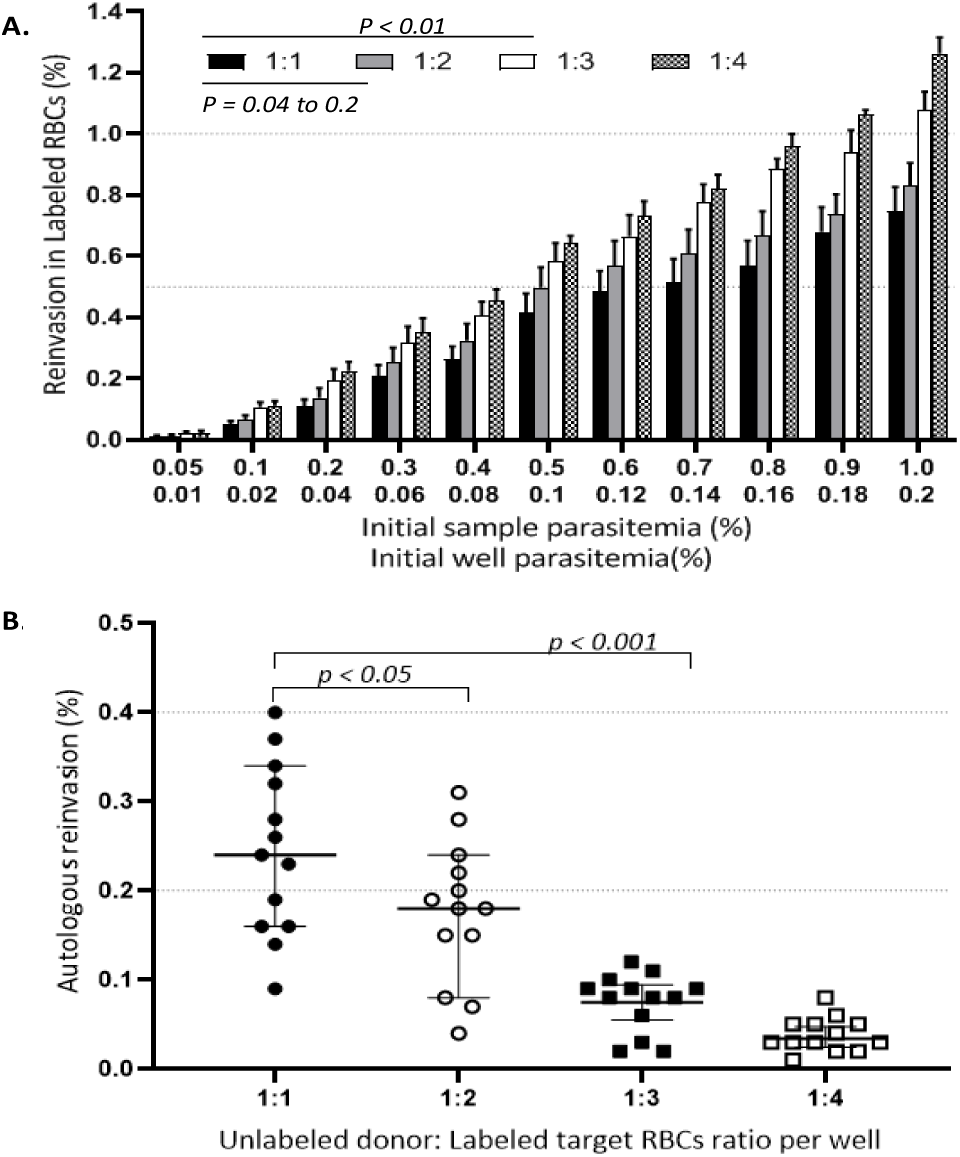
Higher labeled RBCs proportions per assay increases sensitivity of FCM to quantify reinvasion. (A). FCM determined reinvasion for unlabeled to labeled RBCs ratios of 1:1 to 1:4 were compared across separate assays with fixed parasite densities of *Plasmodium falciparum* 3D7 and Dd2 cultures diluted to a parasitemia range of 0.05% to 1.0%. Mean and standard error bars for six replicate values from two experiments are represented. (B). Effect of unlabeled to labeled RBCs ratio on parasites reinvasion into unlabeled uninfected donor RBCs (Autologous reinvasion). 95% CI values of the average of six replicate assays per sample at 0.1% to 1.0% initial parasitemia are represented. Autologous reinvasion (%) **=** (*Final growth %pct* – *Reinvasion %pct*) – (*Initial well %pct*).

### The effect of higher labeled RBCs proportions and initial assay parasitemia on 2cFCM determined invasion pathway phenotypes

Given that reinvasion into unlabeled RBCs was skewed with 1:3 assays (p<0.001), it was next determined if 1:3 assays would improve FCM determination of parasite invasion pathway phenotypes. At this ratio of labeled to unlabeled RBCs, 2cFCM was accurate in recapitulating invasion pathway phenotypes of 3D7 and Dd2 at as low as 0.025% pct per well (0.1% sample parasitemia). This produced a 50% increase in assay sensitivity compared to the conventional 1:1 assay where 2cFCM determination of phenotype was limited to 0.05% pct per well (0.2% sample parasitemia) (figure 5A). *Plasmodium falciparum* Dd2 strain maintained a stable invasion inhibition of >50% by Neuraminidase while the 3D7 strain was inhibited at <50%. This confirmed the respective sialic acid-dependent and independent invasion pathways of these isolates at the new assay cell ratio of 1:3.

**Figure 5.**
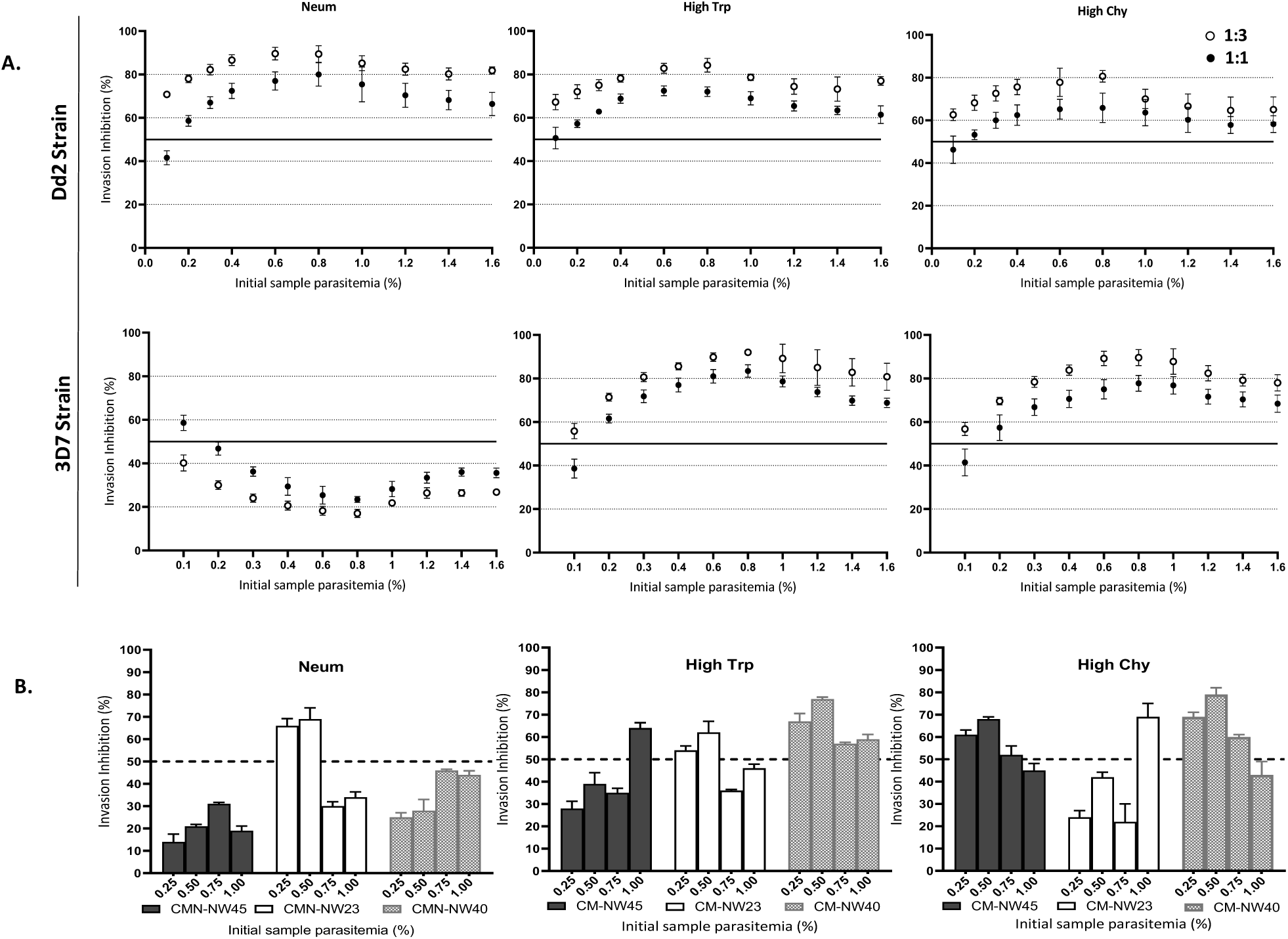
Effect of parasite density and labeled RBCs proportions per assay on FCM determination of invasion pathway phenotypes. (A). Invasion inhibition by enzyme treatments for 1:3 assays (clear squares) and 1:1 assays (dark squares) with Dd2 (upper panel) and 3D7 (lower panel) parasites at an initial sample parasitemia range of 0.1% to 1.6%. (B). Effect of starting parasite density on invasion inhibition by enzyme treatments for samples of field isolates with a 1:3 assay. Labeled target erythrocytes were treated with 3.35mU/ml Neuraminidase, 1.0mg/ml Trypsin and 1.0mg/ml Chymotrypsin. Each point represents the mean + SEM for 2 replicate experiments with triplicate wells per sample. 

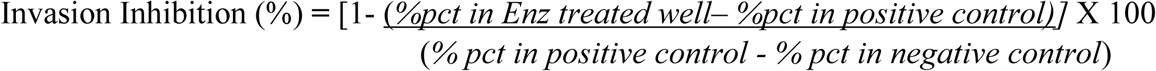

where positive control was Nm+Trp+Chy treated and negative control was non treated RBCs.

Invasion inhibition by enzyme treatment significantly increased by a minimum of 10% across the entire parasitemia range for the 1:3 assay when compared to the 1:1 assay (p<0.05) (figure 5A). Patterns of invasion inhibition were consistent across the sample parasitemia range of 0.1% to 1.6%. There was an increasing trend in the inhibitory effect of enzyme treatments up to an initial sample parasitemia of 0.8%, after which the trend started reversing. This sets 0.8 parasitemia as the peak point of reliable invasion phenotyping for any strain. Surprisingly, RBC invasion by Dd2 strain used in these assays was inhibited by chymotrypsin treatment across the entire parasitemia range tested (figure 5A iii). This contrary results could be due to a reversal in phenotypes that occurs with long-term culture adaptation (Duffy and Avery 2018).

The effect of initial parasitemia on invasion pathways of natural non-culture adapted *P. falciparum* isolates was also investigated on 3 clinical isolates at 0.25%, 0.5%, 0.75% and 1.0% initial parasitemia with the 1:3 protocol. Invasion phenotypes were less consistent across the different initial parasitemia for all enzyme treatments, but invasion phenotypes were similar at parasitemia ≤ 0.5% across all enzyme treatments (figure 5C).

## DISCUSSION

Two-color FCM is the most widely used method for phenotyping parasite invasion pathways as it offers minimal handling of donor infected culture. However, current protocols require samples of field isolates at ∼1% parasitemia for assay set-up (Bei and Duraisingh 2015), which are more uncommon in low transmission settings. This study presents a modification of current protocols to enable the determination of invasion pathways for field isolates at low parasitemia with improved sensitivity and reproducibility.

SYBR Green I has a low binding affinity for RNA and so, will preferentially bind DNA given the shortest time possible. This study protocol reported a reduced parasite DNA staining time with SYBR Green 1 down to 20minutes, from 1 hour used in the conventional protocols (Theron et al. 2010; Bei and Duraisingh 2015). The limit of quantification of reinvasion parasitemia was accurate to 0.05% parasitemia from a 1:1 assay of 0.02% well parasitemia (>2-fold increase after a single cycle of growth). Most studies of invasion pathways reported to date have only assayed isolates with parasitemia within the range of 0.5 to 1% (Bowyer et al. 2015; Mensah-Brown et al. 2015). This reduced staining time saves time to perform assays and minimizes background fluorescence from young unRBCs in which RNA may have not been completely degraded. It equally excludes the need for laborious RNase treatment of RBCs that have been previously explored (Jun et al. 2012; Izumiyama et al. 2009), thereby reducing the cost of invasion phenotyping assays. Increasing the sensitivity of reinvasion quantification at low parasitemia was achieved here by increasing the proportion of labeled target RBCs to two-fold per assay. This resulted in driving increased reinvasion into labeled RBCs (heterologous reinvasion) and consequently minimizing phenotype masking associated with increased autologous reinvasion in 1:1 assays. This suggests that the availability of labeled RBCs in previous protocols was limiting heterologous reinvasion. Earlier modifications of the 2cFCM protocol only compared 1:1 and 1:2 cell ratios in assays (Theron et al. 2010) and did not find labeled RBC availability to be limiting. This new ratio is recommended for adoption as it allows for accurate and sensitive determination of known invasion phenotypes of *P. falciparum* 3D7 and Dd2 strains assayed at 0.1% parasitemia. The increased sensitivity was further enhanced by gating double-positive RBCs in dot plots of SYBY Green I versus CTFR, as a percentage of labeled target RBCs population exclusively. This gating strategy corrects for errors in donor/target RBCs volumes arising from pipetting errors or due to stochastic passing of RBCs via the flow cell which may alter the FCM determination of labeled RBC parasitemia.

However, this revised protocol with a 1:3 ratio of unlabeled to labeled RBCs will result in a higher cost per sample but this can be minimized by setting-up assays in duplicates rather than triplicates. Also, the 1:3 assay further dilutes the initial sample parasitemia by three-fold when compared to one-fold in the 1:1 assay which reduces the sample’s number of parasites. Nonetheless, the impartial invasion of just about all of the parasites sampled per well into labeled RBCs with increased reproducibility and accuracy compensates for the partial invasion of the more sample’s number of per assay well in the 1:1 protocol. Our findings also suggest that high parasitemia like the 0.75% to 1.0% recommended in previous reports (Theron et al. 2010; Bei and Duraisingh 2015) saturates the re-invasion process. Assays at lower parasitemia determined here could be more sensitive, distinguishing smaller differences in invasion phenotypes. This was further demonstrated with 3 field isolates from Cameroon, which are tested for invasion phenotypes for the very first time. As these were complex infections, the effect of clonality and parasite density on invasion pathway phenotypes at these low parasitemia warrant further investigation.

## Conclusion

We have established a revised protocol for the determination of invasion phenotypes of low parasitemia malaria samples. Assays should be set up with an ideal assay (well) parasitemia of 0.05% to 0.2%. The shorter SYBR Green I staining time of 20 minutes and a two-fold increase in labeled target RBCs improves sensitivity and reproducibility of 2cFCM to determine invasion pathway phenotypes, especially at low parasitemia.

## ACKNOWLEDGEMENTS

The authors thank Simon Correa for donating his blood cells for parasite culture and assisting with microscopy readings. We equally thank the malaria patients for donating blood and the staff of the health facilities at the Ndop Health district in the North West region of Cameroon who assisted in patient recruitment. We thank Prof. Cho-Ngwa Fidelis of the University of Buea for providing the dry shipper in which samples were transported to MRC Gambia and the MPB team at MRC Gambia at LSHTM for helpful discussions and advice. We thank Prof Umberto D’Alessandro for supporting the fellowship.

## AUTHORS’ CONTRIBUTIONS

NIA and AAN conceived and designed the experiments, NIA acquired and analyzed data and drafted the manuscript. AAN, FJ, FB and HM made substantial contributions to editing the manuscript. All authors read and approved the final manuscript.

## LIST OF ABBREVIATIONS

FCM: Flow cytometry
RBCs: Red Blood Cells
iRBCs: infected RBCs unRBCs uninfected RBCs
RPMI: Rose Park Memorial Institute iRPMI incomplete RPMI
cRPMI: complete RPMI CTFR Cell Trace Far Red
% pct: Percent parasitemia
Nm: Neuraminidase
Trp: Trypsin
Chy: Chymotrypsin
HCT: Hematocrit

